# ATG4D loss leads to late-onset cardiomyopathy and stress-induced heart failure in mice, and its repression marks maladaptive cardiac remodeling in humans

**DOI:** 10.1101/2025.09.29.678527

**Authors:** Guillermo Mariño, Isaac Tamargo-Gómez, Raquel García-López, Gemma G. Martínez-García, María F. Suárez, Oliva C. Fernández Cimadevilla, Xurde M. Caravia, Raúl F. Pérez, Verónica Rey, María Calvo, J. Francisco Nistal, Álvaro F. Fernández

**Affiliations:** Departamento de Biología Funcional, Facultad de Medicina, Universidad de Oviedo, 33006 Oviedo, Spain; Instituto de Investigación Sanitaria del Principado de Asturias (ISPA), 33011, Oviedo, Spain; Instituto Universitario de Oncología (IUOPA), 33007 Oviedo, Spain; Instituto de Investigación Valdecilla (IDIVAL), 39011 Santander, Spain; Departamento de Fisiología y Farmacología, Facultad de Medicina, Universidad de Cantabria, 39011 Santander, Spain; Servicio de Cardiología, Hospital San Agustín, Avilés, Asturias, Spain; Departamento de Bioquímica y Biología Molecular, Facultad de Medicina, Universidad de Oviedo, 33006 Oviedo, Spain; Departamento de Bioquímica y Biología Molecular, Facultad de Veterinaria, Universidad Complutense de Madrid, 28040 Madrid, Spain; Departamento de Ciencias Médicas y Quirúrgicas, Facultad de Medicina, Universidad de Cantabria, 39011 Santander, Spain; Servicio de Cirugía Cardiovascular, Hospital Universitario Marqués de Valdecilla, 39008 Santander, Spain

## Abstract

In the last years, autophagy has emerged as an essential pathway for most cellular functions. Basal autophagy plays a protective role as a quality control mechanism by which damaged or noxious cellular components are degraded and cellular organelles are periodically renewed. Moreover, autophagic activity can be increased in situations of cellular stress, including nutrient or growth factor deprivation, hypoxia, reactive oxygen species, DNA damage, or the presence of intracellular pathogens. Normally, induction of autophagy is protective, although in some circumstances, such as conditions of hemodynamic stress, autophagosome accumulation upon autophagy induction can be a maladaptive process. The deficiency of the autophagic protease ATG4D in mice leads to the accumulation of cellular autophagosomes in most tissues, including the heart. Here, we show that the increased autophagosome content of *atg4d^−/−^* mice is linked to the development of late-onset cardiomyopathy and to increased susceptibility to heart failure induced by transverse aortic constriction. Furthermore, we report the existence of human *ATG4D* variants associated with cardiovascular pathologies and also that *ATG4D* expression is reduced in human obstructive hypertrophic cardiomyopathy and dilated cardiomyopathy, which highlights a conserved cardio-protective role of the ATG4D protease.

## INTRODUCTION

Cardiovascular diseases (CVDs) are currently the main cause of death at a global scale and, particularly, among the non-communicable diseases^1^. Most of the CVDs-attributable deaths occur in people older than 65 years, which reinforces the concept of CVDs as predominantly aging-related diseases^2^. Healthy aging is characterized by a progressive decrease in both left ventricle (LV) mass and Left Ventricular End-Diastolic Volume (LVEDV)^3^. Conversely, in pathological situations such as CVDs, the LV undergoes a remodeling process that leads to a progressive increase in its mass, usually associated with cardiac hypertrophy^4^. From a cellular point of view, CVDs lead to cardiomyocyte loss and to an increase in extracellular matrix volume triggered by a deregulation of cardiac homeostasis^5^. The maintenance of cardiac health depends on a wide variety of factors, including the activity of the autophagy process, which plays an important role in both cardiac development and cardiac health preservation^6^.

Autophagy is an evolutionarily-conserved catabolic process present in all eukaryotic organisms, from yeast to mammals^7^. During autophagy, different portions of cytoplasm, including organelles, proteins and macromolecules, are sequestered by double-membrane vesicles (called autophagosomes), which eventually fuse with acidic organelles. After this fusion, both the autophagosomal content and the inner autophagosomal membrane are degraded by acidic hydrolases and the resulting products (amino-acids, lipids or nucleotides) are recycled back to the cytosol, where they can be used to generate new cellular structures^8^. Multiple autophagy related (ATG) proteins are required for the correct development of the process^9^. In yeast, one of the essential components for the elongation and expansion of nascent autophagosomes is the Atg4/Atg8 ubiquitin-like conjugation system, in which cytoplasmic protein Atg8 becomes active upon proteolytic cleavage by the cysteine proteinase Atg4^10^. After this priming, active Atg8s are conjugated with phosphatidylethanolamine (PE) molecules from the nascent pre-autophagosomal membrane, allowing the correct formation, closure, and maturation of autophagosomes. Once Atg8 is not required at the autophagosomal membranes, Atg4 is also responsible for its delipidation, allowing its incorporation into newly formed autophagic structures. Contrasting to other autophagy-essential genes, the Atg4/Atg8 ubiquitin-like conjugation system has gained complexity throughout evolution. In fact, there are four human orthologues of Atg4 protease in mammalian cells (ATG4A-D), and six Atg8 orthologues (mATG8s) have been described in humans, organized in two different subfamilies named as LC3 and GABARAP^11^. Although the specific functions of each ATG4 family member are not fully understood, ATG4B protease is the main priming enzyme for mATG8s^12^, whereas ATG4D is the main mATG8s delipidating enzyme in mammalian cells^13^. In turn, ATG4A and ATG4C functions seem to be complementary and redundant to those of ATG4B and ATG4D^14^. Due to its ability to degrade large cellular structures, autophagy contributes to maintaining cellular homeostasis by periodically renewing cytosolic content. Moreover, autophagic activity can be boosted in response to a wide variety of stimuli, preserving cells and tissues health during stressful conditions, such as the accumulation of misfolded proteins, the occurrence of bacterial or viral infections, the presence of damaged organelles, the exposure to environmental pollutants, unhealthy diets, or lack of nutrients by starvation, among many others^15–17^. Although autophagy induction is mostly protective, the exact role of the autophagy pathway in cardiac homeostasis and physiology is controversial. Autophagic activity is necessary for correct heart development and function^18^, and its induction in cardiomyocytes is reported to be beneficial in diverse pathological contexts^19–22^. By contrast, many reports have shown that autophagy induction may be detrimental in the context of CVDs, more explicitly during pressure overload-induced cardiomyopathy^23–28^. Specifically, the excessive activation of the autophagy pathway has been repeatedly reported to play a maladaptive role in pressure overload-induced heart failure. In mice, transverse aortic constriction (TAC) is an established model of elevated afterload, as occurs in the prevalent conditions of hypertension or aortic stenosis. TAC induces tissue remodeling, which is characterized by an increase in the diameter of individual myocytes, accumulation of intercellular collagen (fibrosis), a general increase in the heart muscle mass, dilation of the left ventricle, and functional impairment with a deterioration in the systolic function that may culminate in heart failure and death. Interestingly, hearts from mice subjected to TAC protocols show a massive increase in autophagosome content, which has been repeatedly observed^29, 30^. Far from being just an epiphenomenon accompanying cardiac malfunction, it appears that the accumulation of autophagosomes after TAC is responsible for some of the detrimental effects of pressure overload on cardiac health. In fact, mice haploinsufficient for Beclin-1 (*Becn1*^+/-^) fail to upregulate cardiac autophagy after TAC and are protected from all the manifestations of dilated cardiomyopathy present in banded wild-type (WT) mice^28^. Conversely, transgene-enforced overexpression of Beclin-1 in the heart accelerates TAC-induced heart failure^25^. Moreover, inhibition of autophagosome biogenesis through dimethyl-alpha-ketoglutarate treatment reduces the negative impact of pressure overload and preserves cardiac function in mice subjected to TAC protocols^26, 27^. These observations strongly support the notion that autophagy (or perhaps just autophagosome accumulation) contributes to pathogenic remodeling of the heart muscle under hemodynamic stress.

In this regard, we have recently generated mice deficient for ATG4D, whose study has enabled us to reveal the role of this protease as the main ATG4 member in charge of LC3 and GABARAP proteins delipidation in mammalian cells^31^. Interestingly, *atg4d^−/−^* mice show increased cellular levels of autophagic structures, but no autophagic flux alterations, neither under control conditions nor upon starvation^13^. This prompted us to characterize cardiac function in ATG4D-deficient mice to determine whether the observed increased presence of autophagic structures could, by itself, impact cardiac homeostasis. In this work, we show that ATG4D loss leads to age-associated cardiac hypertrophy and fibrosis, which is preceded by CVD-related features, and confers increased susceptibility to TAC-induced heart failure. In parallel, analysis of data from public repositories revealed the existence of human CVDs-associated *ATG4D* pathological polymorphisms. Consistently, analyses of public transcriptomic datasets revealed a significant downregulation of *ATG4D* expression in myocardial tissue from patients with dilated cardiomyopathy. Moreover, our own analyses identified a similar repression of *ATG4D* expression in interventricular septum samples from patients with obstructive hypertrophic cardiomyopathy (oHCM), a condition which accounts for 10% of sudden cardiac deaths in young persons and a larger proportion in young athletes^32^. Together, these data suggest that ATG4D repression is a conserved feature of pathological cardiac remodeling, which supports a protective role for ATG4D against cardiac dysfunction.

## RESULTS

### *atg4d^−/−^* mice show late-onset cardiomyopathy

It has been proposed that autophagosome accumulation, due to either an increase in their biogenesis or to a block in their degradation, represents a maladaptive response to hemodynamic stress^23, 24, 28^. *atg4d^−/−^* mice show a unique autophagic phenotype characterized by an increased number of autophagosomes in cells and tissues without noticeable alteration of autophagic flux^31^. Interestingly, analysis of publicly available resources from The Human Protein Atlas (https://www.proteinatlas.org) revealed that ATG4D is the most expressed ATG4 protease in the human cardiac left ventricle (Fig. 1A). These results were confirmed in mice using real time qPCR analyses with samples from WT mice left ventricles (Fig. 1B), which could indicate a preponderant role for ATG4D in the heart. Prompted by these premises, we decided to perform a series of analyses focused on the characterization of cardiac physiology in *atg4d^−/−^*mice. We first analyzed the expression of the other members of the ATG4 family in the absence of ATG4D in heart tissue. As shown in (Fig. 1C), both *Atg4c* and *Atg4b* expression are upregulated in *atg4d^−/−^* mice heart tissue, as compared to the levels observed in WT mice, which probably reflects an attempt to compensate for ATG4D absence in this tissue. However, the increase in the expression of *Atg4c* and *Atg4b* is not sufficient to mitigate the consequences of ATG4D loss, especially regarding the delipidation of mATG8 proteins (Fig. 1D-E). In fact, *atg4d^−/−^* mice hearts show increased lipidation of all mATG8 proteins when analyzed by immunoblotting, as previously reported^31^. Although significant for all mATG8 proteins, this observed increase in lipidation is more evident for LC3 proteins and less marked for GABARAP, GABARAPL1 and GABARAPL2 (Fig. 1D-E). Moreover, immunofluorescence analyses of heart tissue sections revealed an increase in the number of positive puncta for each mATG8 protein (corresponding to autophagosomes or autophagosome-related structures) in mutant mice samples when compared to the corresponding WT controls (Fig. 1F-G). Thus, given the observed alterations in the status of mATG8 proteins in the hearts of ATG4D-deficient mice, and due to the importance of autophagy for cardiac physiology, we decided to perform histopathological and functional analyses in the hearts of *atg4d^−/−^* mice at different ages. As shown in Figure 2, knockout mice do not exhibit any evident cardiac alteration at a young age (2 months), neither as measured by gross anatomical and histopathological analyses (Fig. 2A-C), nor using echocardiography (ECHO) (Fig. 2D-E, S1A). However, when we extended the ECHO-based analyses to middle-aged mice (9 to 12 months), we were able to observe a clear cardiac phenotype in *atg4d^−/−^* mice, which exhibit signs of eccentric hypertrophy, with increased LV dimensions, both systolic and diastolic, and a reduced relative posterior wall thickness (Fig. 2F-G, S1B). These more spherical LVs maintained preserved systolic (in both axes) and diastolic functions, as evidenced by MAPSE values similar to those of WT littermates, although we could observe a trend pointing to an age-dependent reduction in the ejection fraction and LV fractional shortening of mutant mice that did not reach statistical significance, likely due to variability in the data (Fig. 2G, S1B). In this sense, post-mortem histopathological analyses in older mice (18 to 26 months old) revealed an increased incidence of features associated with hypertrophic cardiomyopathy, such as increased heart size, higher heart/body mass ratio and left ventricular dilatation as compared with their WT littermates (Fig. 2H-J). Consistently, wheat germ agglutinin (WGA) staining revealed a significant increase in the area occupied by connective tissue and also in the average cardiomyocyte size, usually a sign of cardiac cardiomyocyte hypertrophy (Fig. 2K). These alterations were more frequent in female mice and were associated with a higher incidence of myocardial fibrosis, as shown by trichromic staining (Fig. 2L) and confirmed by TEM analyses (Fig. 2M). Interestingly, although the overall survival of ATG4D-deficient mice is not statistically different from that of WT mice, when sex-dependent long-term survival was analyzed, we could observe that female *atg4d^−/−^* mice showed a significant reduction in their average and maximal lifespan (Fig. 2N). This reduction in female *atg4d^−/−^* mice survival could be linked to the observed cardiac hypertrophy and fibrosis, which could predispose them to premature death.

**Figure 1.**
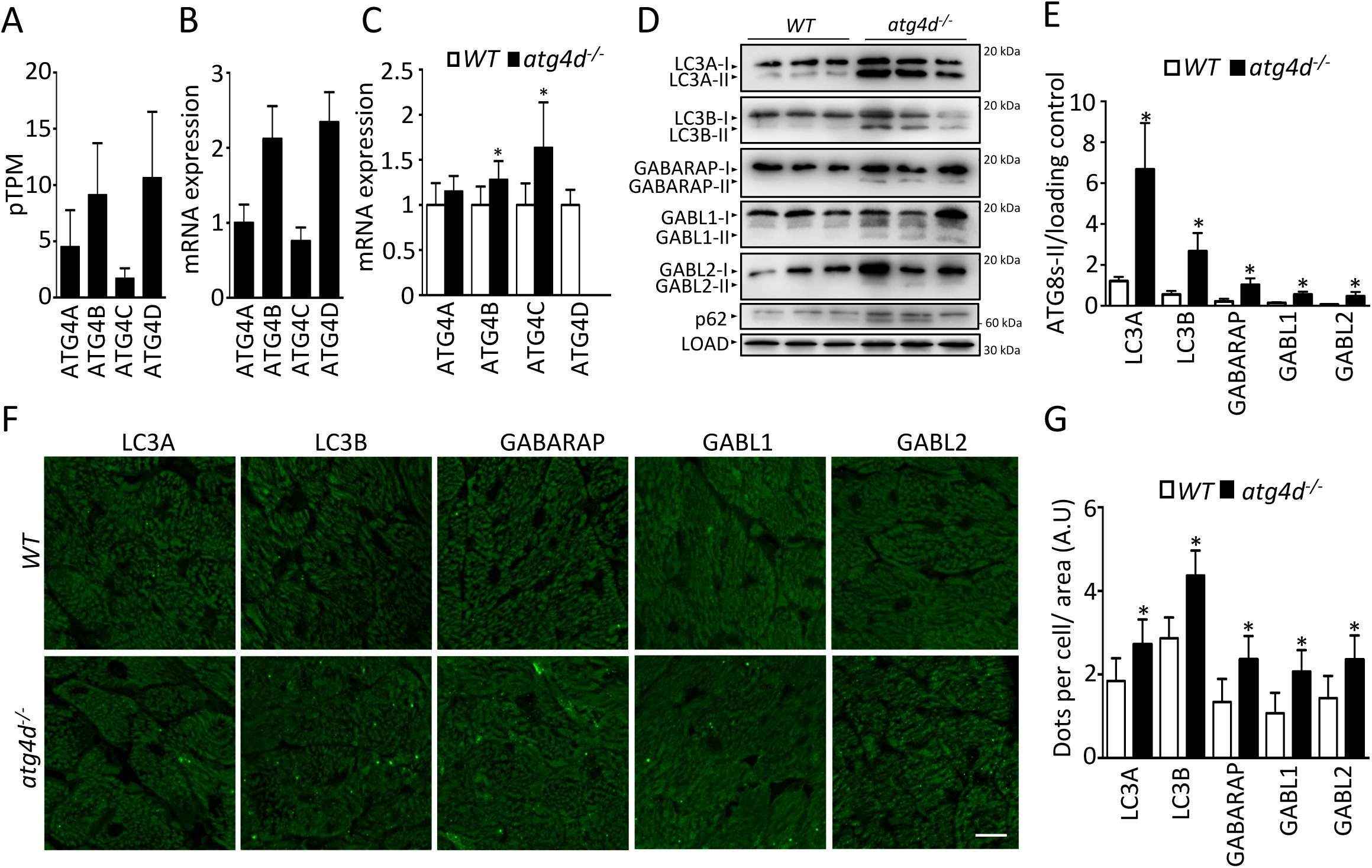
mATG8 status in WT and *atg4d*^−/−^ mice hearts. (A) pTPM expression of each ATG4 protein in the left ventricle of human samples. n = 303 samples; data obtained from https://www.proteinatlas.org. (B) mRNA expression of different ATG4s in the left ventricle of 2-month-old WT mice samples. Values were relativized to ATG4A expression. n = 6 mice. (C) mRNA expression of ATG4D in the left ventricle of 4-month-old WT and *atg4d*^−/−^ mice. (D) Representative immunoblots of endogenous ATG8-like proteins in the hearts from 2-month-old WT and *atg4d*^−/−^ mice, showing an increase of their lipidated forms in mutant mice. (E) Quantification of immunoblots in (D). (F) Representative images of immunofluorescence analysis of endogenous mATG8 proteins in WT and *atg4d*^−/−^ hearts. (G) Quantification of the data from (F). n = 4 mice per genotype. Bars represent mean ± SEM; *, p < 0.05, 2-tailed unpaired Student’s t test. n = 3 mice per genotype.

**Figure 2.**
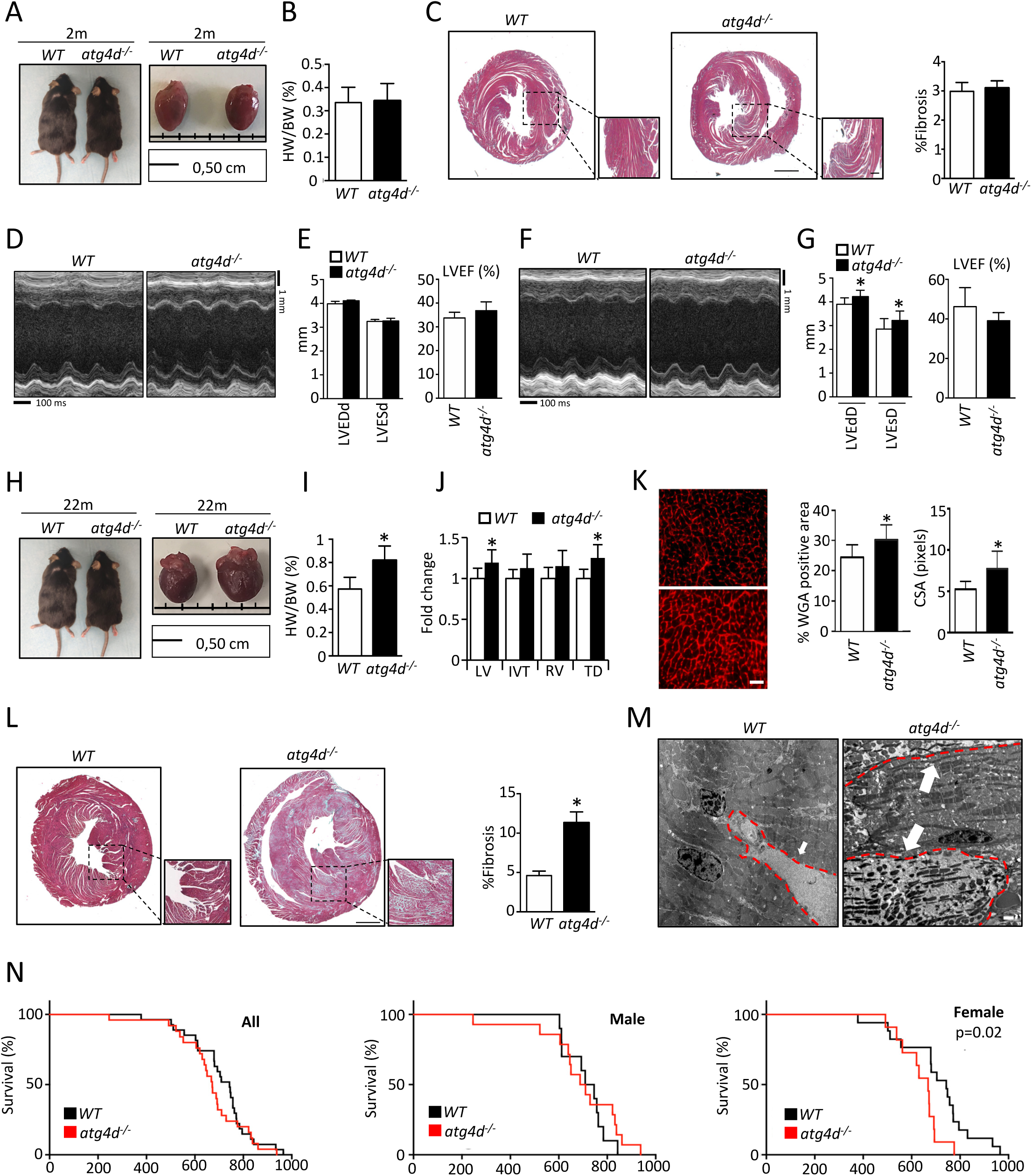
Age-dependent cardiac remodeling and reduced survival in *atg4d*^−/−^ mice. (A) Representative images of mice and excised hearts. (B) Evaluation of heart weight/body weight (HW/BW) ratio in 2-month-old WT and *atg4d*^−/−^ mice. n = 7 mice per genotype. (C) Left, trichromic staining in heart sections from 2-month-old WT and *atg4d*^−/−^ mice. Scale bar: 50 μm, 8 μm in insets. Right, quantification of interstitial fibrosis in 2-month-old WT and *atg4d*^−/−^ mice. n = 5 mice per genotype. (D) Representative M-mode echocardiographic images in 2-month-old WT and *atg4d*^−/−^ mice. (E). Quantification of left ventricular end-diastolic (LVEDd) and end-systolic (LVESd) diameters (left) and left ventricular ejection fraction (LVEF) (right) in 2-month-old WT and *atg4d*^−/−^ mice. (F). Representative M-mode echocardiographic images in 10-month-old WT and *atg4d*^−/−^ mice. (G). Quantification of left ventricular end-diastolic (LVEDd) and end-systolic (LVESd) diameters (left) and left ventricular ejection fraction (LVEF) (right) in 10-month-old WT and *atg4d*^−/−^ mice. (H). Representative images of mice and excised hearts. (I) Evaluation of heart weight/body weight (HW/BW) ratio in 22-month-old WT and *atg4d*^−/−^ mice. (J). Quantitative assessment of left and right ventricular chamber dimensions, interventricular septum thickness, and overall cardiac diameter in 22-month-old WT and *atg4d*^−/−^ mice. Values are expressed relative to those of WT controls. n = 5 mice per genotype. (K) Left, wheat germ agglutinin (WGA) staining in left ventricular cross sections from 22-month-old WT and *atg4d*^−/−^ mice. Scale bar: 20 μm. Right, quantification of cardiomyocyte cross-sectional area and percentage of WGA-positive area. n = 8 mice per genotype. (L) Left, trichromic staining in heart sections from 22-22-month-old WT and *atg4d*^−/−^ mice. Scale bar: 50 μm, 8 μm in inserts. Right, quantification of fibrosis (in percentage). n = 8 mice per genotype. (M) Representative transmission electron microscopy (TEM) images showing the ultrastructure of 22-month-old WT and *atg4d*^−/−^ mice myocardia. White arrows show areas of fibrosis (between sarcomeres and perivascular in WT and replacing degenerated sarcomeres in *atg4d*^−/−^ samples). Scale bar, 4μm. (N) Kaplan–Meier survival curves for WT and *atg4d*^−/−^mice showing the lifespan of all mice in the cohort (left), males alone (middle) and females alone (right). (n = 27 WT and 25 KO mice). In all graphs, bars represent mean ± Standard deviation. p-values were determined by a 2-tailed unpaired Student’s t-test. In (N), p-values were determined by the log-rank (Mantel–Cox) test. *, p < 0.05.

### *atg4d^−/−^* mice show reduced cardiac fitness in response to hemodynamic stress

Cardiac overload due to elevated blood pressure has been reported to induce myocardial autophagosome accumulation, which most likely compromises autophagosome clearance due to insufficient lysosomal content and interferes with cardiac function^33^.

Thus, in order to better characterize the role of ATG4D in the heart, we decided to evaluate the susceptibility of *atg4d^−/−^*mice to experimentally induced hypertension. To this aim, we first subjected WT and *Atg4d*-null mice to chronic Angiotensin-II infusion, which leads to cardiac remodeling as a consequence of pressure overload^34–37^. After one week of Angiotensin-II treatment, *atg4d^−/−^* mice showed a comparable incidence of infarctions and fibrotic lesions to those observed in their WT counterparts (Fig. 3A-C). However, lesions from *Atg4d*-null mice showed increased necrosis severity and a worse clinical score than WT animals (Fig. 3D-E). These results suggest that ATG4D deficiency leads to a reduced adaptability of heart tissue in this experimental setting.

**Figure 3.**
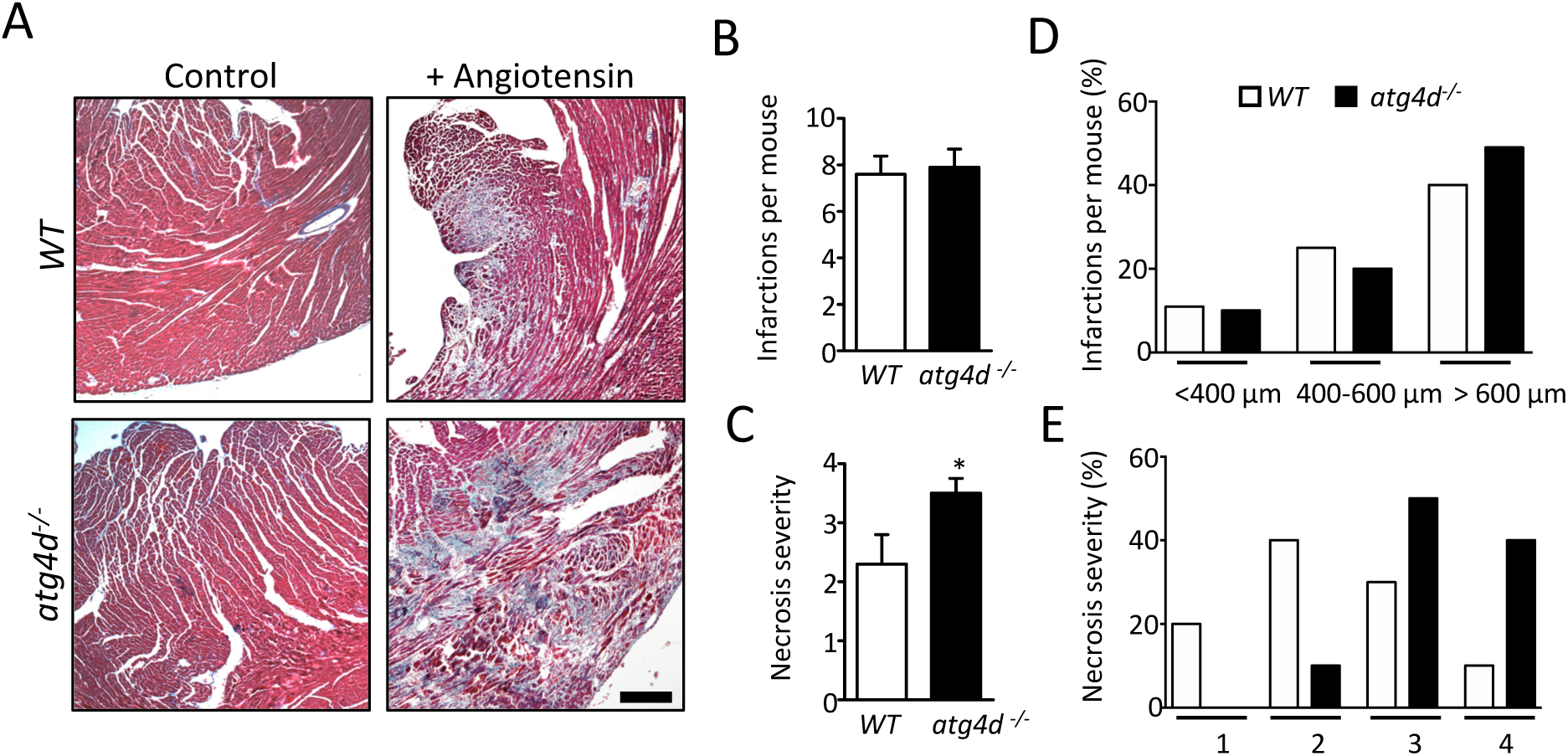
Analysis of Angiotensin II treatment in 2-month-old WT and *atg4d^−/−^* mice. (A) Representative pictures of trichromic staining in heart sections from angiotensin II-treated mice. For a week, 2-month-old WT and *atg4d*^−/−^ mice were treated with Angiotensin II, which leads to cardiac remodeling and hypertrophy because of blood pressure overload. (B) Number of infarctions per mouse in 2-month-old WT and *atg4d*^−/−^ mice subjected to Angiotensin II-induced experimental hypertension. (C) necrosis severity of the infarctions. (D) Distribution of infarcts according to the size of the lesion. < 400 µm, 400-600 µm and > 600 µm. (E) Distribution of infarcts according to grade of necrosis suffered. Grade 1 to 4. Scale bar: 50 μm. Bars represent mean ± SEM *, p < 0.05, 2-tailed unpaired Student’s t test.

Prompted by these findings, and to further analyze the roles of ATG4D in heart physiology, we subjected 2- and 9-month-old WT and *Atg4d*-null mice to controlled transverse aortic constriction (TAC), which is a commonly used experimental model for pressure overload-induced cardiac hypertrophy and heart failure^38^. As expected, TAC led to the accumulation of lipidated forms of all mATG8s in WT mice, as detected by immunoanalysis of heart tissue extracts (Fig. 4A-B). This increase in mATG8 lipidation was particularly evident for LC3A, LC3B and GABARAPL2 and less marked for GABARAP and GABARAPL1 proteins (Fig. 4A-B). Interestingly, although *atg4d^−/−^* mice samples already show accumulation of lipidated forms of all mATG8 proteins at basal conditions, we could observe a further increase in the lipidation of most of them upon TAC (Fig. 4A-B). These results were confirmed using immunofluorescence analyses of all five endogenous mATG8 proteins, as we observed an increase in the number of positive structures for these proteins (mostly indicative of autophagosomes) in heart sections from WT mice subjected to TAC (Fig. 4C-D). In fact, *atg4d^−/−^*mice hearts showed an increased number of mATG8-positive structures when compared to WT animals, either at basal conditions or upon TAC (Fig. 4C-D). These analyses confirm that knockout mice are able to increase their cardiac autophagosome content further upon TAC, despite their already higher autophagosome number at basal conditions. To extend these analyses, we differentially compared the lipidation status of mATG8 proteins between the right and left ventricles from WT and mutant mice subjected to TAC (Fig. S2A-B). Interestingly, *Atg4d*-deficient mice showed a significant increase in the lipidation of all mATG8 proteins in right ventricular extracts as compared to those from *WT* mice (Fig. S2A-B). However, the increase in mATG8s lipidation in *atg4d^−/−^* mice samples was restricted to LC3B, GABARAP and GABARAPL2 when the left ventricle alone was analyzed (Fig. S2A-B). Irrespective of these considerations, ECHO-based evaluation of morphological, geometric and functional changes in the hearts of WT and *atg4d^−/−^* mice 8 and 19 days after surgery showed interesting results, revealing an increased susceptibility of mutant mice to experimentally-induced hemodynamic stress. At a young age (2-month-old), *atg4d^−/−^* mice already showed signs of increased sensitivity to TAC-induced cardiac overload, although our results did not reach statistical significance (Fig. S3). However, when older, 10-12 months old mutant mice that were subjected to TAC exhibited greater LV dilatation at end-diastole and end-systole than their WT counterparts, a more significant time progression of IVS thickening, greater LV mass growth, more deterioration of short-axis systolic function, and a higher level of transcoarctational gradient across TAC (Fig. 5A-B, S4). Moreover, the echocardiographic functional differences between WT and *atg4d^−/−^*mice subjected to TAC were associated with a higher presence of cardiac fibrosis in mutant mice, as shown by Gomori trichromic staining (Fig. 5C-D). Furthermore, WGA staining revealed a significant increase in cross-sectional area, which points to increased cardiomyocyte hypertrophy in *atg4d*^−/−^ mice (Fig. 5E-F).

**Figure 4.**
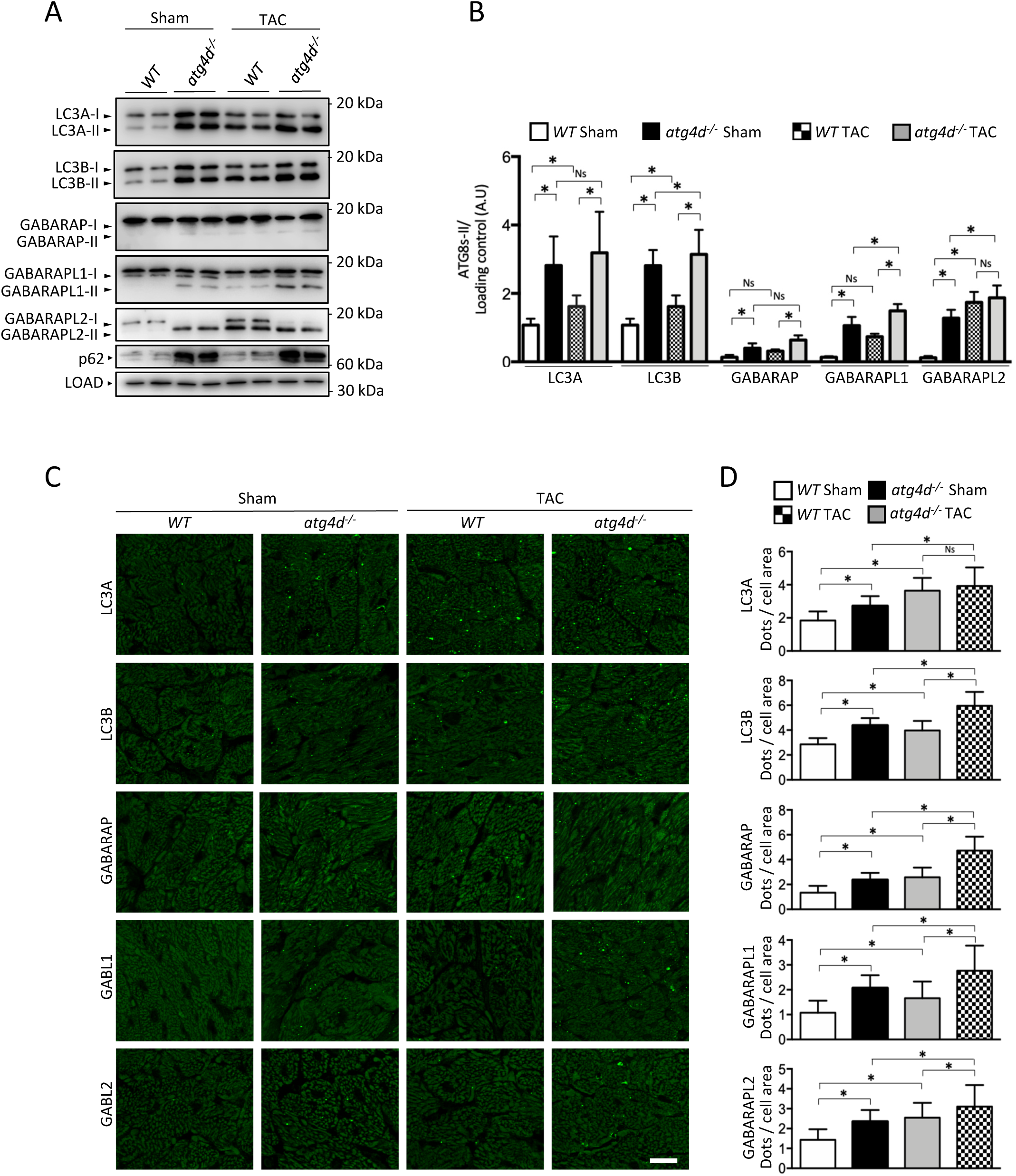
Analysis of ATG8-like proteins in WT and *atg4d*^−/−^ mice upon transverse aortic constriction (TAC). (A) Representative immunoblots of endogenous ATG8-like proteins in hearts from control WT and knock-out mice and in WT and *atg4d*^−/−^ hearts upon TAC treatment, showing an increase of their lipidated forms and an increase in the levels of the autophagy adaptor SQSTM1/p62 in mutant mice. (B) Quantification of immunoblots in (A). (C) Representative images of immunofluorescence analysis of endogenous mATG8 proteins in WT and *atg4d*^−/−^ hearts in control and upon TAC conditions. (D) Quantification of the data from (A) n = 4 mice per condition and genotype. Scale bar: 20 μm. Bars represent mean ± SEM *, p < 0.05, two-way ANOVA test.

**Figure 5:**
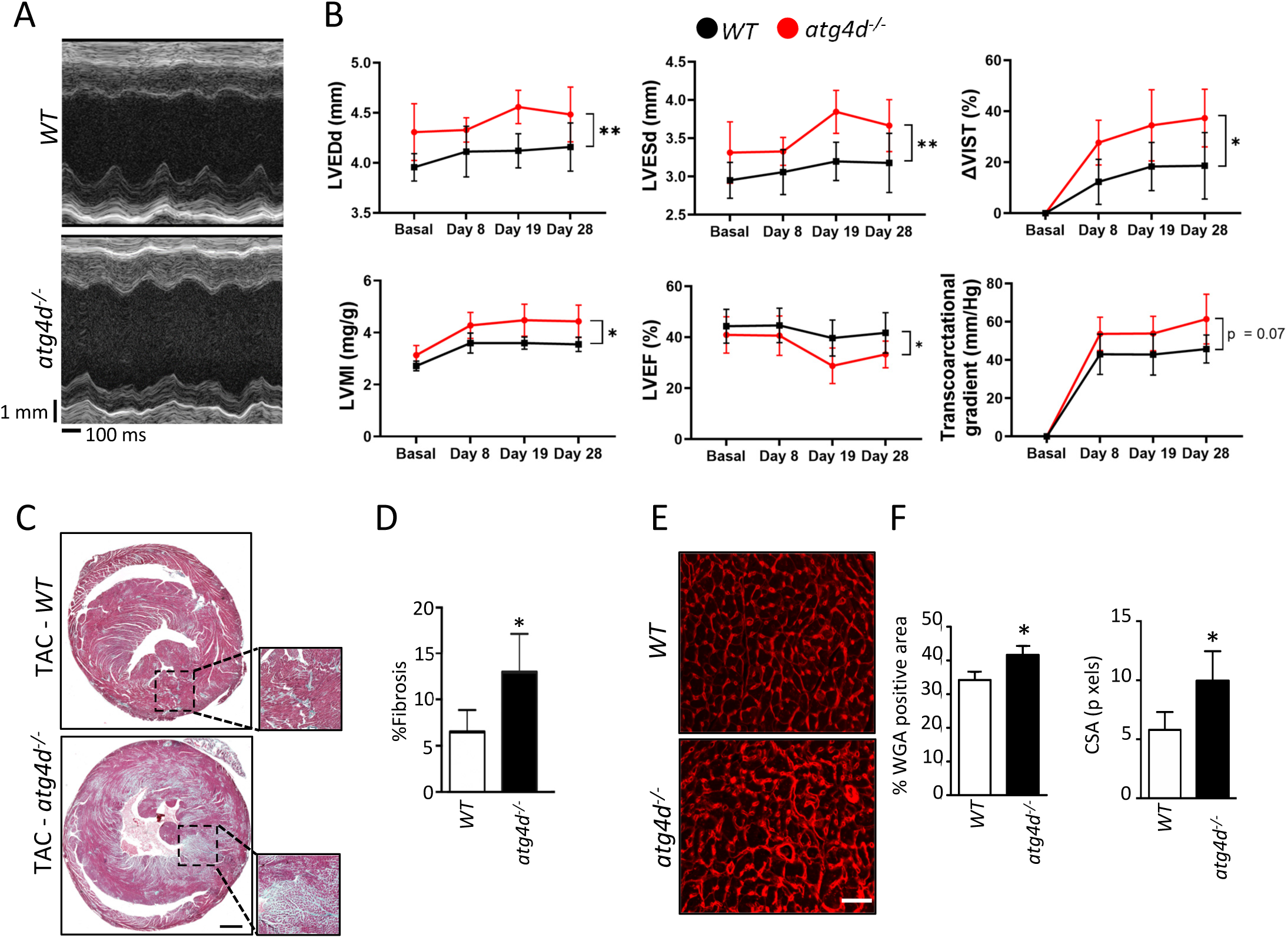
Exacerbated cardiac remodeling in *atg4d*^−/−^ mice subjected to 4-week transverse aortic constriction (TAC). (A) Representative M-mode echocardiographic images of 10-month-old WT and *atg4d*^−/−^ mice at day 28 after cardiac stress induction. (B) Time-course analysis of echocardiographic parameters: left ventricular end-diastolic diameter (LVEDd), end-systolic diameter (LVESd), interventricular septal thickening (ΔIVST), left ventricular mass (LVM), ejection fraction (LVEF), and transaortic pressure gradient in 10-month-old WT and *atg4d*^−/−^ mice during TAC. Bars represent mean ± standard deviation *p < 0.05, in repeated measures two-way ANOVA (C) Representative heart sections stained with Masson’s trichrome in 10-month-old WT and *atg4d*^−/−^ mice at day 28 after TAC. Scale bar: 50 μm, 8 μm in inserts. (D) Quantification of fibrosis represented in (C). (n = 4 WT, n = 7 KO). (E) Wheat germ agglutinin (WGA) staining in left ventricles transverse sections from WT and *atg4d*^−/−^ mice at day 28 after TAC. Scale bar: 20 μm (F) Quantification of cardiomyocyte cross-sectional area and percentage of WGA-positive area. in D and F, bars represent mean ± SEM *, p < 0.05, 2-tailed unpaired Student’s t test.

Together, these results show that the absence of ATG4D in mice increases their susceptibility to both Angiotensin-II infusion and surgical transverse aortic constriction, increasing heart hypertrophy and fibrosis in response to hemodynamic stress. All these features are associated with an increased lipidation of mATG8 proteins and with the accumulation of autophagic structures, which may be the underlying cause of the increased susceptibility of *atg4d*^−/−^ cardiomyocytes to the detrimental effects of experimentally induced cardiac overload.

### Single-nucleotide polymorphisms (SNPs) and expression levels of *ATG4D* are associated with cardiovascular function

The fact that *atg4d*^−/−^ mice develop age-associated cardiomyopathy together with their increased susceptibility to both Angiotensin-II infusion and transverse aortic constriction prompted us to perform an exhaustive search in publicly accessible databases (https://cvd.hugeamp.org; https://www.ebi.ac.uk/gwas/) to find putative single-nucleotide polymorphisms (SNPs) in *ATG4D* and in other different autophagy-related genes, which could be associated with cardiovascular function. As shown in Table 1, we could find a high number of SNPs in the *ATG4D* gene, which were associated with different alterations in cardiac function. Specifically, several *ATG4D*-related variants are associated with ischemic pathologies, alterations in left ventricular end-systolic volume, and myocardial infarctions, while some of them are related to changes in LDL cholesterol. Interestingly, *ATG4D* showed the highest number of SNPs associated with cardiovascular pathologies among all autophagy-related genes, although we could also find a variety of SNPs in several other ATG genes, which were linked to this type of condition (Table 1). Transcriptomic evidence from both experimental and clinical sources also reinforces the notion that ATG4D plays a crucial role in maintaining cardiac homeostasis and that its downregulation may be a consistent feature of pathological cardiac remodeling. In WT mice subjected to TAC, *Atg4d* expression was found to be significantly downregulated when compared to sham-operated controls (Fig. 6A). This observation is supported by reanalysis of publicly available gene expression data from TAC-operated mice^39^ (GSE56348), which includes over 40 autophagy-related genes (Table S1). Among them, *Atg4d* showed one of the strongest and most statistically significant reductions in expression (Fig. 6B-C), suggesting that suppression of this gene may be a specific and early molecular signature of the maladaptive response to sustained hemodynamic stress. The transcriptomic downregulation of different autophagy genes under these conditions may reflect impaired activation or progression of the autophagic program, yet the particularly prominent decrease in *Atg4d* levels highlights this gene as a key node potentially disrupted during disease onset. Importantly, this pattern of *Atg4d* repression is not restricted to animal models. Analysis of myocardial tissue derived from patients with clinically diagnosed dilated cardiomyopathy (DCM) also revealed a marked reduction in *ATG4D* expression compared to non-failing human hearts^40^. In this dataset (GSE26887), which encompasses a large panel of autophagy-related transcripts, *ATG4D* emerged as the most significantly downregulated gene within the autophagy machinery (Fig. 6D-E and Table S1), further supporting the clinical relevance of its suppression. Finally, to directly explore whether *ATG4D* dysregulation also occurs in human hypertrophic remodeling, we analyzed LV posterior wall samples from patients undergoing septal myectomy for obstructive hypertrophic cardiomyopathy (oHCM) and compared them with non-LV pressure overloaded surgical controls. Quantitative RT–PCR revealed a significant reduction of *ATG4D* mRNA in oHCM myocardium as compared with controls (Fig. 6F). The consistency of these findings across species and disease models indicates that the loss of *ATG4D* expression may be linked to progressive cardiac dysfunction.

**Figure 6:**
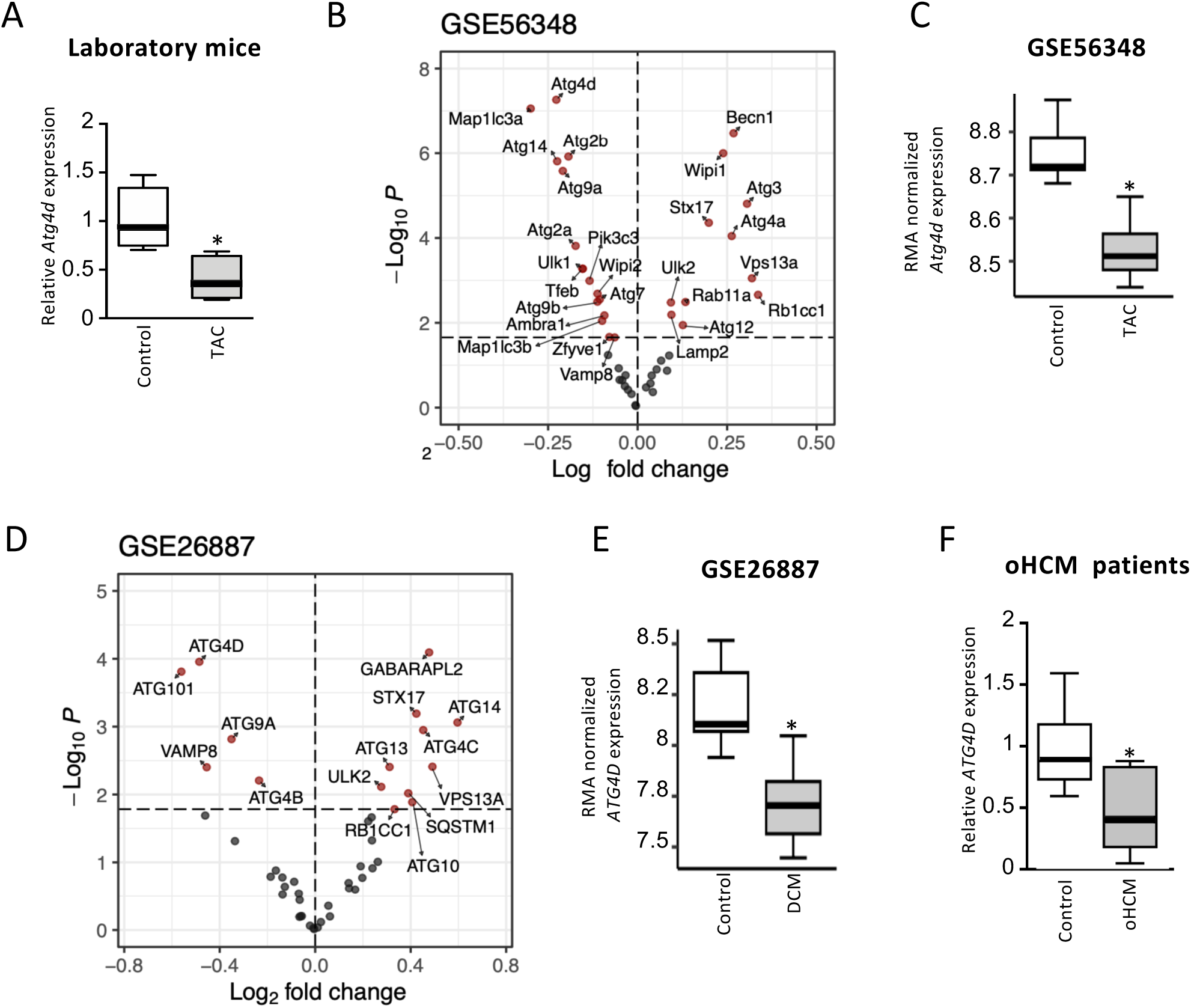
ATG4D is downregulated during heart failure (HF). (A) Real-time PCR analysis showing relative expression of *Atg4d* in left ventricles from 2-month-old *Wild-type* mice subjected to TAC. n = 5 mice per condition. Volcano plot showing the autophagy-related genes significantly altered between heart failure mouse models and controls from Lai L. et al^39^ (dataset GSE56348). Genes with FDR<0.05 are colored in red. C) Boxplot highlighting the specific differences observed for the *Atg4d* gene in GSE56348 dataset. D) Volcano plot showing the autophagy-related genes significantly altered between heart failure human subjects and controls from Greco S. et al^40^ (dataset GSE26887). Genes with FDR<0.05 are colored in red. E) Boxplot highlighting the specific differences observed for the *ATG4D* gene in the GSE26887 dataset. F) Boxplot representation of relative ATG4D mRNA levels in LV posterior wall samples from non-pressure overloaded human hearts (control) and patients with oHCM undergoing septal myectomy. Data are shown as box-and-whisker plots (min–max) with median values indicated. *, p < 0.05, 2-tailed unpaired Student’s *t* test.

**Table 1.** Analysis of *ATG4D* polymorphisms related to cardiovascular function. Representative SNPs of *ATG4D* associated with different cardiovascular functions. Data obtained from https://hugeamp.org and https://www.ebi.ac.uk/gwas/.

| Variant ID | dbSNP ID | Major allele | Minor allele | Sequence ontology<br>TERM | Biological relation | Gene |
| --- | --- | --- | --- | --- | --- | --- |
| 19:10663568:GCTC:G |  | GCTC | - | inframe_deletion | Myocardial infarction | *ATG4D* |
| 19:10655489:G:A | rs200278873 | G | A | missense_variant | Ischemic stroke | *ATG4D* |
| 19:10663165:C:T | rs7255312 | C | T | intron_variant | Ischemic stroke | *ATG4D* |
| 19:10659659:C:T | rs2304165 | C | T | synonymous_variant | Ischemic stroke | *ATG4D* |
| 19:10662236:T:C | rs2011172 | T | C | intron_variant | Left ventricular end-systolic volume | *ATG4D* |
| 19:10663997:C:T | rs77956256 | C | T | 3_prime_UTR_variant | Left ventricular end-systolic volume | *ATG4D* |
| 19:10663568:G:GCTC |  | G | GCTC | inframe_insertion | Left ventricular end-systolic volume | *ATG4D* |
| 19:10660685:C:A | rs139718563 | C | A | intron_variant | Left ventricular end-systolic volume | *ATG4D* |
| 19:10654705:C:T | rs10423320 | C | T | 5_prime_UTR_variant | Coronary artery disease | *ATG4D* |
| 19:10658257:T:C | rs62128796 | T | C | intron_variant | Coronary artery disease | *ATG4D* |
| 19:10663165:C:T | rs7255312 | C | T | intron_variant | Coronary artery disease | *ATG4D* |
| 19:10548983:C:T | rs2304165 | C | T | synonymous_variant | Coronary artery disease | *ATG4D* |
| 19:10551099:T:C | rs8108051 | T | C | intron_variant | Coronary artery disease | *ATG4D* |
| 19:10549564:T:C | rs8103305 | T | C | intron_variant | Coronary artery disease | *ATG4D* |
| 19:10550594:C:T | rs58265604 | C | T | upstream_gene_variant | Coronary artery disease | *ATG4D* |
| 19:10542342:G:A | rs12459544 | G | A | upstream_gene_variant | Coronary artery disease | *ATG4D* |
| 19:10550944:T:A | rs12973308 | T | A | intron_variant | Coronary artery disease | *ATG4D* |
| 19:10541931:C:T | rs10408372 | C | T | upstream_gene_variant | Coronary artery disease | *ATG4D* |
| 19:10659659:C:T | rs2304165 | C | T | missense_variant | LDL cholesterol | *ATG4D* |
| 19:10654705:C:T | rs10423320 | C | T | 5_prime_UTR_variant | LDL cholesterol | *ATG4D* |
| 19:10658257:T:C | rs62128796 | T | C | intron_variant | LDL cholesterol | *ATG4D* |
| 19:10659737:C:T | rs12979378 | C | T | intron_variant | LDL cholesterol | *ATG4D* |
| 19:10645495:A:G | rs10439163 | A | G | intergenic_variant | LDL cholesterol | *ATG4D* |
| 19:10549574:C:T | rs35041134 | C | T | intron_variant | LDL cholesterol | *ATG4D* |
| 19:10549382:C:A | rs34479588 | C | A | intron_variant | LDL cholesterol | *ATG4D* |
| 19:10542528:G:A | rs111428249 | G | A | upstream_gene_variant | LDL cholesterol | *ATG4D* |
| 19:10542744:A:G | rs73496510 | A | G | upstream_gene_variant | LDL cholesterol | *ATG4D* |
| 19:10551335:G:A | rs111573781 | G | A | intron_variant | LDL cholesterol | *ATG4D* |
| 19:10550523:C:T | rs73496517 | C | T | intron_variant | LDL cholesterol | *ATG4D* |
| 19:10543267:C:T | rs73496512 | C | T | upstream_gene_variant | LDL cholesterol | *ATG4D* |
| 19:10550773:A:C | rs35643447 | A | C | intron_variant | LDL cholesterol | *ATG4D* |
| 19:10550944:T:G | rs12973308 | T | G | intron_variant | LDL cholesterol | *ATG4D* |
| 19:10542394:G:C | rs12459547 | G | C | upstream_gene_variant | LDL cholesterol | *ATG4D* |
| 19:10547759:C:T | rs75120078 | C | T | intron_variant | LDL cholesterol | *ATG4D* |
| 19:10549466:G:A | rs12460639 | G | A | intron_variant | LDL cholesterol | *ATG4D* |
| 19:10550509:T:C | rs11666992 | T | C | intron_variant | LDL cholesterol | *ATG4D* |
| 19:10551392:A:G | rs10411361 | A | G | intron_variant | LDL cholesterol | *ATG4D* |
| 19:10546352:A:T | rs113370024 | A | T | intron_variant | LDL cholesterol | *ATG4D* |
| 19:10552103:G:A | rs148200963 | G | A | synonymous_variant | LDL cholesterol | *ATG4D* |
| 19:10549059:A:G | rs3816042 | A | G | intron_variant | LDL cholesterol | *ATG4D* |

Taken together, the downregulation of ATG4D observed both in murine models of pressure overload and in human DCM and HOCM hearts, as well as the correlation between a high number of *ATG4D* SNPs and the development of cardiovascular pathologies, point to a cardio-protective role for *ATG4D* protease in mice and humans. Further investigation into the regulatory mechanisms governing *ATG4D* transcription in the heart, as well as the functional consequences of its loss in vivo, will be critical to understanding its role in cardiac pathophysiology.

## DISCUSSION

Autophagy is an essential pathway for most cellular functions, either as a basal housekeeping route or as a pro-survival process against a variety of intracellular or extracellular stressors. Due to its complexity, the autophagy route implies the function of more than 30 autophagy-specific genes/proteins (ATG) in mammalian cells^41^. In the last years, the generation of animal models deficient for ATG genes has boosted our knowledge about the implications of the autophagy process for cell, tissue and organismal homeostasis^42^.

Even though autophagic activity is low in cardiomyocytes under basal conditions, it acts as a protective mechanism against the accumulation of toxic protein aggregates in specific situations of cardiac stress^43^. In fact, a deficiency of autophagy core genes causes important alterations in cardiac function. In this sense, *Atg5* loss in cardiomyocytes leads to heart failure in mice, which is characterized by reduced ejection fraction, hypertrophy, fibrosis, and an early decline in cardiac function with a premature mortality in approximately 10 months^44^. Conversely, autophagy induction can be maladaptive for cardiomyocytes in some particular circumstances. This is illustrated by the improved response of autophagy-haploinsufficient *Becn1*^+/-^ mice to hemodynamic stress^28^. Specifically, *Becn1*^+/-^ mice are protected from the detrimental effects of experimentally induced pressure overload. Interestingly, a reduced content of autophagic structures is generated in response to TAC in these mice, as compared with WT animals. On the other hand, genetically increasing the rate of autophagosome biogenesis by Beclin-1 overexpression leads to the development of more severe clinical features in mice subjected to the same experimental settings^23^. A similar situation is observed in the case of familial Danon disease patients, who develop fatal cardiomyopathy, among other clinical features. Danon disease is caused by mutations in the *LAMP2* gene, which ultimately lead to autophagosome accumulation in cardiomyocytes as a consequence of autophagosome-lysosome fusion impairment^45^. Mice deficient for LAMP2 phenocopy human Danon disease, showing autophagosome accumulation in cardiomyocytes, which ultimately leads to severe attenuation of cardiac contractile function, among other features^46^. Consistently, samples of septal myectomies from patients suffering from hypertrophic cardiomyopathy show accumulation of early and late autophagic vacuoles and higher protein levels of LC3B-II and Beclin-1 compared with donor samples^47^.

Among autophagy-related proteins, ATG4D plays a specific and crucial role in the delipidation of LC3 and its paralogs from autophagic membranes, a process essential for the recycling and maturation of autophagosomes. Unlike other ATG4 family members, ATG4D has been shown to regulate autophagic turnover by ensuring the correct balance between lipidated and delipidated pools of mATG8s^48^. We have previously shown that the absence of ATG4D leads to an abnormal accumulation of autophagic vesicles both in cultured cells and in mouse tissues^48^. Consistently, the results presented in this work confirm that the loss of ATG4D in cardiomyocytes leads to increased lipidation of mATG8 proteins and an accumulation of autophagic structures. This accumulation of autophagosomes, despite an otherwise intact autophagic flux^48^, suggests an imbalance in the dynamic equilibrium of autophagic membrane turnover. As cardiac tissue relies on a tightly regulated proteostatic network to sustain contractile function, the persistence of excess autophagosomes may impose a burden on intracellular trafficking, mitochondrial maintenance, and sarcomeric integrity. Over time, this could result in a decline in myocardial performance and cardiac adaptability, predisposing to the development of age-related cardiomyopathies and increasing susceptibility to pathological stimuli, such as pressure overload, which increases metabolic demand and mechanical stress (Fig. 7). Thus, our results strongly support the idea that increased autophagosome content in cardiomyocytes (even when it is not derived from a block in autophagosome degradation) compromises cardiac function. Importantly, the existence of human *ATG4D* variants associated with cardiovascular pathologies, together with our experimental evidence, supports the view that ATG4D acts as a conserved cardio-protective factor. In mice, we demonstrate that *Atg4d* expression is significantly reduced following transverse aortic constriction (TAC), and this finding is consistent with results reported by other laboratories, consolidating the notion that *Atg4d* repression is an early and robust response to pressure overload. In humans, independent transcriptomic studies have shown that *ATG4D* is markedly downregulated in myocardial tissue from patients with dilated cardiomyopathy, further reinforcing the clinical relevance of this pathway. Extending these observations, our own analysis reveals that *ATG4D* expression is also reduced in LV wall samples from patients with obstructive hypertrophic cardiomyopathy (oHCM), underscoring that *ATG4D* repression is a recurrent molecular feature across distinct etiologies of pathological cardiac remodeling. In this regard, autophagy has been found to be altered in several HCM scenarios^49, 50^, and its modulation has been proposed as a potential therapeutic target for the disease^51^. Given ATG4D’s essential role in mATG8 delipidation and autophagosome turnover, its suppression may promote autophagosome accumulation, disrupt proteostatic balance, and compromise myocardial resilience under sustained hemodynamic stress. The convergence of data from murine models and human pathologies therefore strengthens the translational significance of ATG4D and raises the possibility that restoring its activity could mitigate maladaptive remodeling in both hypertrophic and failing hearts. From a pharmacological standpoint, ATG4D represents an attractive target due to its enzymatic nature, and strategies aimed at preserving or enhancing its activity may help maintain cardiac structure and function.

**Figure 7:**
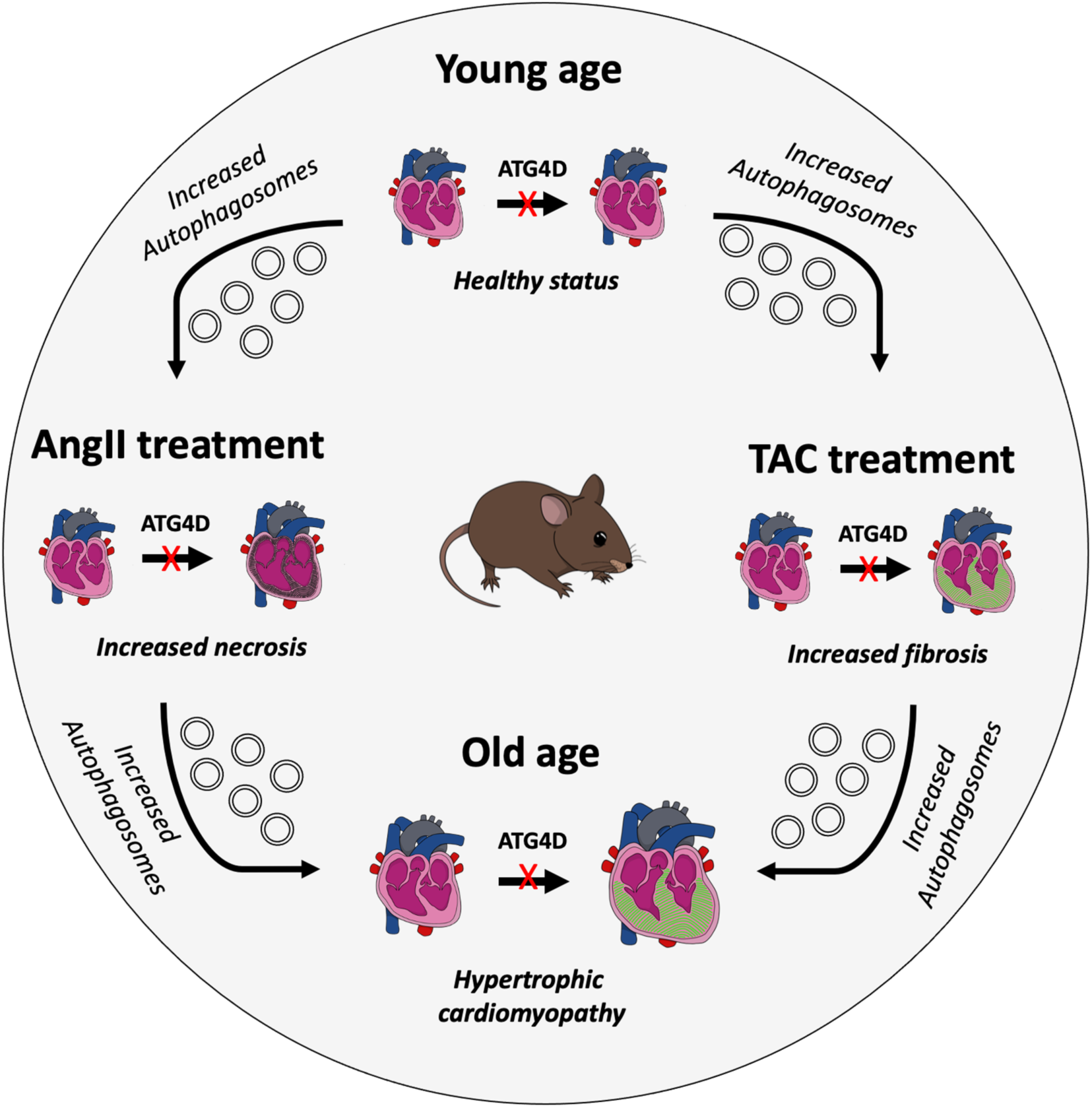
Scheme depicting the consequences of ATG4D loss. At the organismal level, ATG4D loss does not cause any obvious pathology at a young age in unchallenged mice. However, *Atg4d*-deficient mice show late-onset hypertrophic cardiomyopathy, increased susceptibility to Angiotensin II-induced experimental hypertension and morpho-functional echocardiographic changes induced by pressure overload in transverse aortic constriction.

In conclusion, our findings highlight ATG4D as a central regulator of cardiomyocyte autophagy and cardiac physiology, and suggest that its repression is a unifying hallmark of adverse remodeling across species and disease contexts. Further elucidation of the upstream pathways controlling ATG4D expression, as well as the development of pharmacological modulators of its activity, will be critical to establish new therapeutic approaches for cardiovascular disease.

## MATERIALS AND METHODS

### Mice strains

*Atg4d*-knockout mice were generated at the University of Oviedo facilities using the targeted ES cell clones (OST254045) that were obtained from Lexicon Genetics, as previously reported^31^.

### Antibodies

In this study, the following antibodies were used:

Anti-LC3A (Proteintech, Cat# 12135-1-AP); anti-LC3B (Novus, Cat# NB600-1384); anti-LC3C (D3O6P) (Cell Signaling Technology, Cat# 14736); anti-GABARAP (MBL International, Cat# PM037); anti-GABARAPL1 (ATG8L) (Proteintech, Cat# 11010-1-AP); anti-GABARAPL2 (GATE-16) (MBL International, Cat# PM038); anti-GFP (Abcam, Cat# ab290); and anti-GAPDH (Novus Cat# NB300-320).

### Oligonucleotides

In this study, the following oligonucleotides were used:

*Atg4d* mouse genotyping:

5’-CAGACCGCAGGAAAGCAAGGTAT-3’;
5’-AAATGGCGTTACTTAAGCTAGCTTG-3’;
5’-AGTATAGAGTAACACTGTGCTGGC-3’

### Histology analyses

Tissues were fixed in 4% paraformaldehyde in phosphate-buffered saline (PBS) and stored in 50% ethanol. Fixed tissues were embedded in paraffin by standard procedures. Blocks were sectioned (5 μm), and hematoxylin–eosin (HE), Masson’s and Gomori’s Trichrome staining were performed. All tissues were examined by a pathologist in a blinded fashion. For the Angiotensin II–induced cardiac fibrosis protocol, after 7 days of Angiotensin II infusions, hearts were harvested, fixed in paraformaldehyde 4% in PBS, embedded in paraffin, sectioned every 200 μm, and stained with Gomori’s Trichrome. Fibrotic lesions were classified into three categories: Small (< 200 μm), medium (> 200 μm and < 400 μm), and large (> 400 μm), depending on the number of correlative heart sections where lesions appeared. Fibrotic extension was graded from 0 to 4 according to a clinical score. Grade 0: no fibrotic areas; grade 1: less than 25% of myocardium affected; grade 2: from 26% to 50% of myocardium affected; grade 3: from 51% to 75% of myocardium affected; and grade 4: more than 76% of myocardium affected.

### Immunofluorescence analyses

For immunofluorescence analyses with tissue cryo-sections, sections were pretreated for 30 minutes in 1% H2O2/PBS, followed by 1 hour in blocking solution and incubated overnight at 4°C with the correspondent primary antibody (1:100 in PBS). Next day, samples were washed 3 times in PBS for 15 minutes each. Then, sections were incubated with secondary antibody (1:300 in PBS) at RT for 1 hour, thoroughly washed in PBS 3 times, and stained with DAPI for nuclear staining.

### RT-PCR

Total RNA was isolated from mouse tissues according to the method of Chomczynski and Sacchi^52^. About half of the obtained product was reverse-transcribed using the RNA-PCR Core kit® from Perkin-Elmer (Roche Applied Science, Indianapolis, IN). A PCR reaction was then performed with mouse *Atg4d* specific primers for 25 cycles of denaturation (94 °C, 20 seconds), annealing (62 °C, 20 seconds), and extension (72 °C, 30 seconds). As a control, actin was PCR-amplified from all samples under the same conditions.

### Quantitative real-time PCR

cDNA was synthesized using 1 to 5 µg of total RNA, 0.14 mM oligo(dT) (22-mer) primer, 0.2 mM concentrations each of deoxynucleoside triphosphate and SuperScript II reverse transcriptase (Invitrogen). Quantitative RT-PCR (qRT-PCR) was carried out in triplicate for each sample using 20 ng cDNA, TaqMan Universal PCR Master Mix (Applied BioSystems) and 1 μl of the specific TaqMan custom gene expression assay for the gene of interest (Applied Biosystems). To quantify gene expression, PCR was performed at 95°C for 10 minutes, followed by 40 cycles at 95°C for 15 seconds, 60°C for 30 seconds and 72°C for 30 seconds using an ABI Prism 7700 Sequence Detection System. As an internal controlfor the amount of template cDNA used, gene expression was normalized to the mouse β-actin gene using the Mouse β-actin Endogenous Control (VIC/MGB Probe, Primer Limited). Relative expression was calculated as RQ=2^-ΔΔCt^.

### Protein extract preparation

Tissues were immediately frozen in liquid nitrogen after extraction and homogenized in a 20 mM Tris buffer pH 7.4, containing 150 mM NaCl, 1% Triton X-100, 10 mM EDTA and Complete® protease inhibitor cocktail (Roche Applied Science). Then, tissue extracts were centrifuged at 12.000 rpm at 4°C and supernatants were collected. Protein concentration was quantified by bicinchoninic acid technique (BCA protein assay kit, Pierce Biotechnology, 23225). For protein extracts derived from cultured cells, cells were washed with cold PBS and lysed in a buffer containing 1% NP-40, 20 mM HEPES (pH 7.9), 10 mM KCl, 1 mM EDTA, 10% glycerol, 1 mM orthovanadate, 1 mM phenylmethanesulfonyl fluoride (PMSF), 1 mM dithiothreitol, 10 µg/ml aprotinin, 10 µg/ ml leupeptin and 10 µg/ml pepstatin. Lysates were centrifuged at 12.000 rpm at 4°C and supernatants were collected. Protein concentration was quantified by bicinchoninic acid technique (BCA protein assay kit, Pierce Biotechnology, 23225).

### Immunoblotting

A total of 25 µg of protein sample was loaded on either 8% or 13% SDS-polyacrylamide gels. After electrophoresis, gels were electrotransferred onto polyvinylidene difluoride (PVDF) membranes (Millipore), and then membranes were blocked with 5% non-fat dried milk in PBT (phosphate-buffered saline with 0.05% Tween 20) and incubated overnight at 4°C with primary antibodies diluted in 3% non-fat dried milk in PBT. After three washes with PBT, membranes were incubated with the corresponding secondary antibody at a 1:10.000 dilution in 1.5% milk in PBT and were developed with Immobilon Western Chemiluminescent HRP substrate (Millipore, P36599A) by using Odyssey® Fc Imaging System (LI-COR, Lincoln, NE, USA). Unless otherwise specified, immunoblotting against GAPDH was used as a sample processing control (LOAD) for the immunoblots shown in this article.

### Mouse echocardiography

High-resolution transthoracic echocardiography (Visual Sonics Vevo-770, Fujifilm-Visual Sonics, Toronto, Ontario, Canada) was performed in mice under sedation with isoflurane vaporized at 2.5% in oxygen for induction and 1-1.5% for data recording. Care was taken to avoid hypothermia of the animals and, to this end, core temperature was monitored with a rectal probe and active heat was provided through the exploration platform and with an infrared lamp. The minimum heart rate considered acceptable for the studies was 400 beats/minute. Measurements were taken according to the recommendations of the American Society of Echocardiography.

The study protocol included the following projections and measurements:

Right parasternal view of the aortic arch, parasternal long-axis view of the heart, parasternal short-axis view and four-chamber view. These projections allowed the measurement and calculation of: end-diastolic LV diameter (LVEDD), end-systolic LV diameter (LVESD), interventricular septal diastolic thickness (IVS), posterior wall diastolic thickness (PWT), relative posterior wall thickness (rPWT) as a measure of concentric geometry (rPWT=(2xPWT)/LVEDD), LV ejection fraction (Quiñones), mitral annular plane systolic excursion (MAPSE) as a measure of longitudinal LV systolic function, mitral early diastolic peak flow to peak early diastolic myocardial velocity in the mitral annulus (E/e’) as a surrogate of LV filling pressures, left ventricular mass (LVM, Devereux), and transcoarctational pressure gradient (Bernoulli modified formula).

### Transverse aortic constriction (TAC)

Transverse aortic constriction was performed, as previously reported^37^, using a double-loop-clip technique with a 7/0 polypropylene suture (Prolene™ 7/0) with two reference knots. Briefly, the aortic arch was approached, under spontaneous ventilation, through a midline extrapleural route. Once an adequate view of the aortic arch was obtained, the aorta was surrounded, between the origins of the innominate arterial trunk and the left carotid artery, with an atraumatic dissector that allowed the arch to be encircled and the TAC suture to be threaded through twice. Subsequently, both suture ends were gently pulled until the two knots were level, placing a vascular microclip just below them (Weck Horizon microclips, Teleflex Medical, Research Triangle Park, North Carolina). The perioperative mortality of the procedure was 3%.

### Transmission electron microscopy

For ultrastructural studies, cultured cells were fixed in 1.6% glutaraldehyde (v:v in 0.1 M phosphate buffer) for 1 hour, scraped off the plastic dish, centrifuged and post-fixed as a cell pellet in 1% osmium tetroxide (w:v in 0.1 M phosphate buffer) for 2 hours. Following dehydration through a graded ethanol series, cells were embedded in EponTM 812. Ultrathin sections were stained with standard uranyl acetate and lead citrate. For immunogold studies, cells were fixed with either 4% formaldehyde or 1.6% glutaraldehyde in 0.1 M phosphate buffer (pH 7.3) for 1 hour at 4°C. Cell pellets were dehydrated in methanol and embedded in Lowicryl K4M at −20°C in an AFS2 Freeze Substitution Processor apparatus (Leica Microsystems). Polymerization under UV light was carried out for 2 days at −20°C, followed by 2 days at 20°C. Ultrathin sections were incubated with primary antibodies specific to GFP (#ab290, Abcam) for 1 hour at room temperature, and then with secondary antibodies conjugated to 10- or 15-nm gold particles (BBI International, Cardiff, UK), as appropriate. Images were acquired with a Tecnai 12 electron microscope (FEI, Eindhoven, the Netherlands). For electron microscopy of mouse tissues, tissue samples were harvested from mice and immediately fixed in 3% glutaraldehyde in 0.1 M sodium cacodylate (pH 7.2) overnight. After 3 washes in 5% sucrose in 0.1 M sodium cacodylate buffer, samples were post-fixed with 1% osmium tetroxide for 1 hour in the dark and rinsed 3 times in 0.1 M sodium cacodylate. Tissues were dehydrated with increasing acetone concentrations: 30% for 10 minutes, 60% for 10 minutes, 90% for 10 minutes and 100%, three times, for 10 minutes each. The dehydrated pieces were then immersed in mixtures of anhydrous acetone and resin (Durcapan^(R)^ ACM, Fluka BioChemika) of increasing resin concentrations (1:1, 1:2), each step for 30 minutes, and then in pure resin (at 37°C for 12 hours, followed by 60°C for 24 hours). Ultra-thin sections (85 nm) were taken from each sample and analyzed on a Jeol (JEM-1011).

### Obtention of human samples (oHCM)

The study was performed using LV myocardial intraoperative biopsies obtained from surgical patients with pathologies that did not result in LV pressure or volume overload, coronary heart disease or cardiomyopathies (n=7), and a group of patients with obstructive primary hypertrophic cardiomyopathy (oHCM) (n=7). The clinical and demographic characteristics of these groups are shown in Table 2.

**Table 2.**
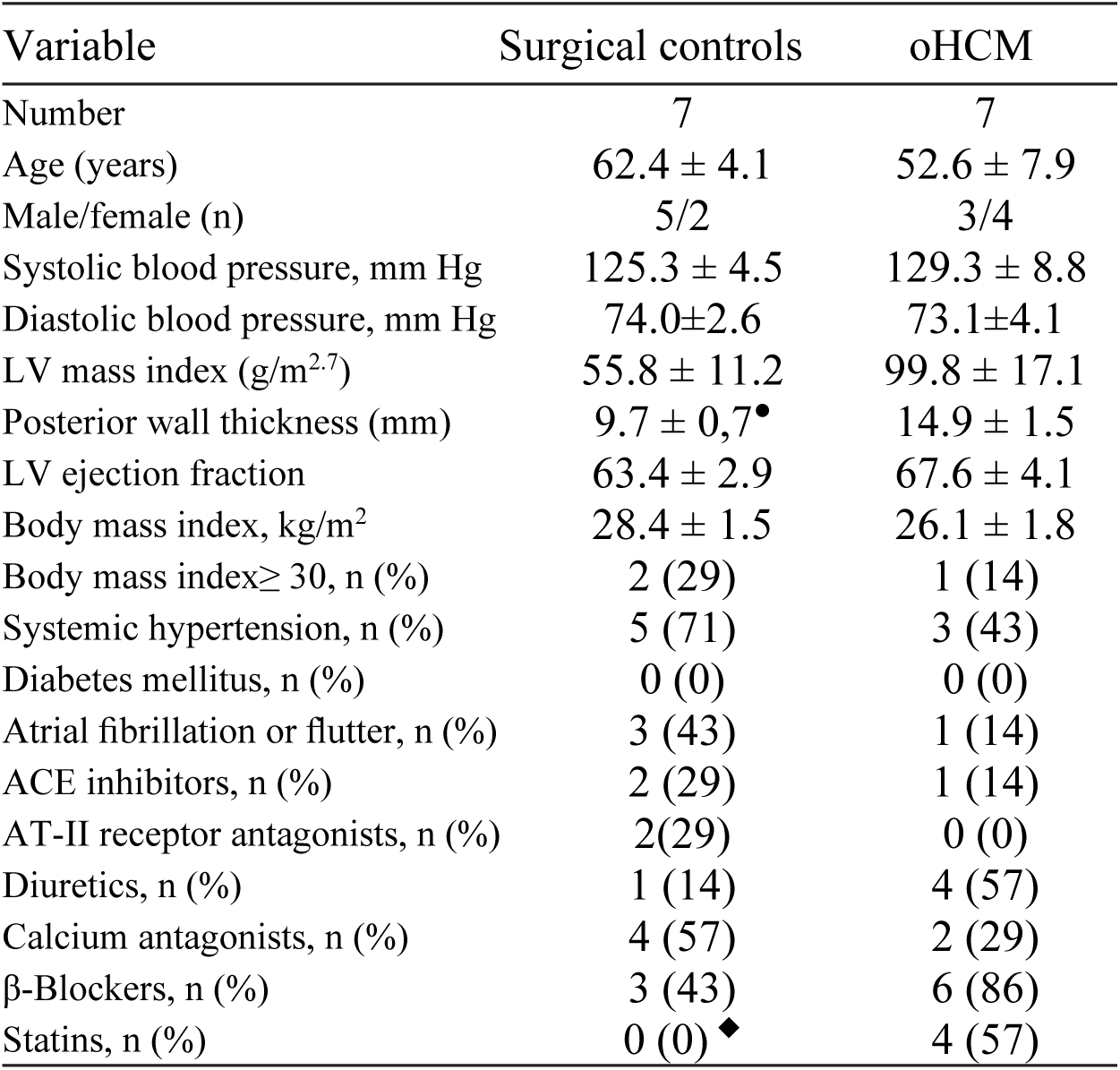
Clinical characteristics of the human study subjects. Patient demographic and clinical continuous variables are presented as mean±standard error of the mean. ACE: Angiotensin converting enzyme; AT-II: Angiotensin-II; LV: left ventricle; oHCM: obstructive hypertrophic cardiomyopathy. ●, p<0.01 vs oHCM (Student’s *t* test); ◆, p<0.05 vs oHCM (Fisher’s exact test).

Prior to surgery, a two-dimensional transthoracic echocardiogram was performed (Philips-Hewlett Packard, IE 33). Left ventricular wall thicknesses were measured according to the American Society of Echocardiography guidelines, using bidimensional or M-mode images depending on the quality and angulation between the ultrasound beam and the LV.

LV mass (LVM) was estimated according to the Devereux formula and indexed to patient height in meters to the 2.7th power. All patients underwent surgery in the University Hospital Marqués de Valdecilla in Santander, Spain. Myocardial tru-cut needle subepicardial biopsies (40 mg) were obtained from the lateral LV wall during surgery and were immediately snap frozen in liquid nitrogen.

### Analysis of external gene expression data

Gene expression data for human and mouse experimental models of cardiomyopathy were downloaded from Gene Expression Omnibus^53^ GSE26887, comprising data of left ventricle biopsies from human patients with heart failure (diabetic subjects were filtered out)^40^; GSE56348, containing data from two mouse models of heart failure and of transverse aortic constriction^39^ (subjects related to exercise models were filtered out). All analyses were performed using R software (v4.3.1). In brief, CEL files were preprocessed via the *oligo* package (v1.66.0)^54^ making use of the appropriate annotation packages for each array: *pd.hugene.1.0.st.v1* (v3.14.1), *hugene10sttranscriptcluster.db* (v8.8.0), *pd.mogene.1.0.st.v1* (v3.14.1), *mogene10sttranscriptcluster.db* (v8.8.0). Next, data were normalized by the RMA method, summarized to the gene level and annotated to genes. Probesets not mapping to any gene, or mapping to >1 different genes were filtered out, while those mapping to the same gene were averaged. Duplicated probesets displaying identical measurements were filtered out. Then a selection of autophagy-related genes was evaluated in the datasets (*ATG10*, *ATG101*, *ATG12*, *ATG13*, *ATG14*, *ATG2A*, *ATG2B*, *ATG3*, *ATG4A*, *ATG4B*, *ATG4C*, *ATG4D*, *ATG5*, *ATG7*, *ATG9A*, *ATG9B*, *ATG16L1*, *ATG16L2, ULK1*, *ULK2*, *MAP1LC3A*, *MAP1LC3B*, *GABARAP*, *GABARAPL1*, *GABARAPL2*, *VAMP8*, *STX17*, *MTOR*, *TFEB*, *UVRAG*, *AMBRA1*, *WIPI1*, *WIPI2*, *RAB11A*, *BECN1*, *LAMP2*, *LAMP1*, *SQSTM1*, *RB1CC1*, *RUBCN*, *NBR1*, *PIK3C3*, *VPS13A*, *VPS13C*, *ZFYVE1*). Differential expression comparisons between phenotype and control groups were carried out with moderated t-tests were used via the *limma* package (v3.58.1)^55^. Graphs were built using *ggplot2* (v3.5.13.3), and *EnhancedVolcano* (v1.20.0).

### Quantification and statistical analysis

All experimental data are reported as mean ± SEM unless otherwise mentioned. Normality of the variables was tested by means of the Shapiro-Wilk test. Unpaired two-tailed Student’s *t*-test was used when comparing two experimental groups, while three experimental groups were analyzed using *one-way ANOVA* followed by Tukey’s post hoc test. Comparison of baseline echocardiographic and left ventricular gene expression data as a function of genotype and pressure overload status was compared by two-way ANOVA.

The influence of age or genotype on the time course of pressure overload-induced modifications in echocardiographic parameters was assessed by a repeated-measures two-way ANOVA. The Bonferroni post-hoc test was used when appropriate. Chi-square was applied for some alignment analysis. Statistical packages: GraphPad Prism 5.03 (Graph-Pad Software Inc., San Diego, CA, USA) and PASW Statistics 18 (SPSS Inc., Chicago, IL, USA). *P* values lower than 0.05 were considered significant.

## ACKNOWLEDGEMENTS

The authors thank A. Vincelle for assistance in characterizing the cardiovascular phenotype, N. García-Iglesias for technical support, and R. Moreta, R.N., together with Beatriz García-Cañón, R.N., for their careful management of clinical samples.

## AUTHOR CONTRIBUTIONS

G.M., I.T-G., X.M.-C., Á.F.F., M.F.S., G.G.M-G. and O.F.C. participated in the generation, analysis and maintenance of *Atg4d*-knockout mice, as well as performed *in vivo* experiments with *Atg4d*-knockout mice; M.F.S. and I.T-G. performed immunoblotting and immunofluorescence experiments; Á.F.F, R. F. P. performed statistical analyses of the expression of autophagy-related genes. I. T-G. performed the search for *ATG4D* SNPs. V.R. helped with histological analyses of hearts; J.F.N., R.G.L., and M.C. performed TAC experiments echocardiographic monitoring and experiments with human samples; G.M. wrote the manuscript, devised the concept and supervised the project.

## COMPLIANCE WITH ETHICAL STANDARDS

Experiments with human samples followed the Declaration of Helsinki guidelines for investigation in humans. The institutional ethics and clinical research committee of the University Hospital Valdecilla approved the study, and all enrolled patients gave written informed consent. Animal experiments in this article were approved by the Committee on Animal Experimentation of Universidad de Oviedo (Oviedo, Spain) (PROAE 10/2022).

