## Supplementary figures and images for "ATG4D loss leads to late-onset cardiomyopathy and stress-induced heart failure in mice, and its repression marks maladaptive cardiac remodeling in humans"

### Supplementary figures

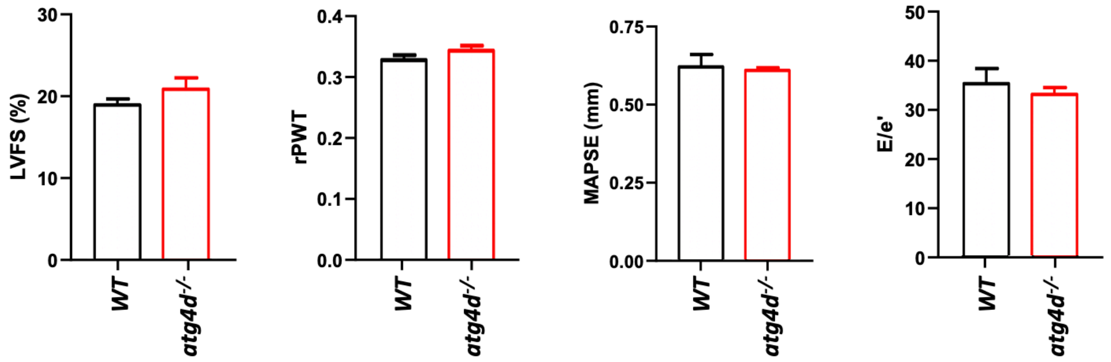

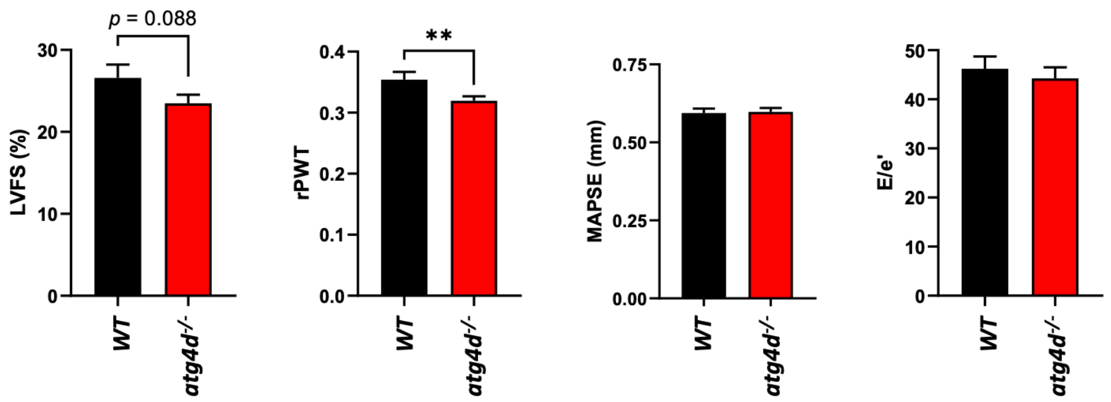

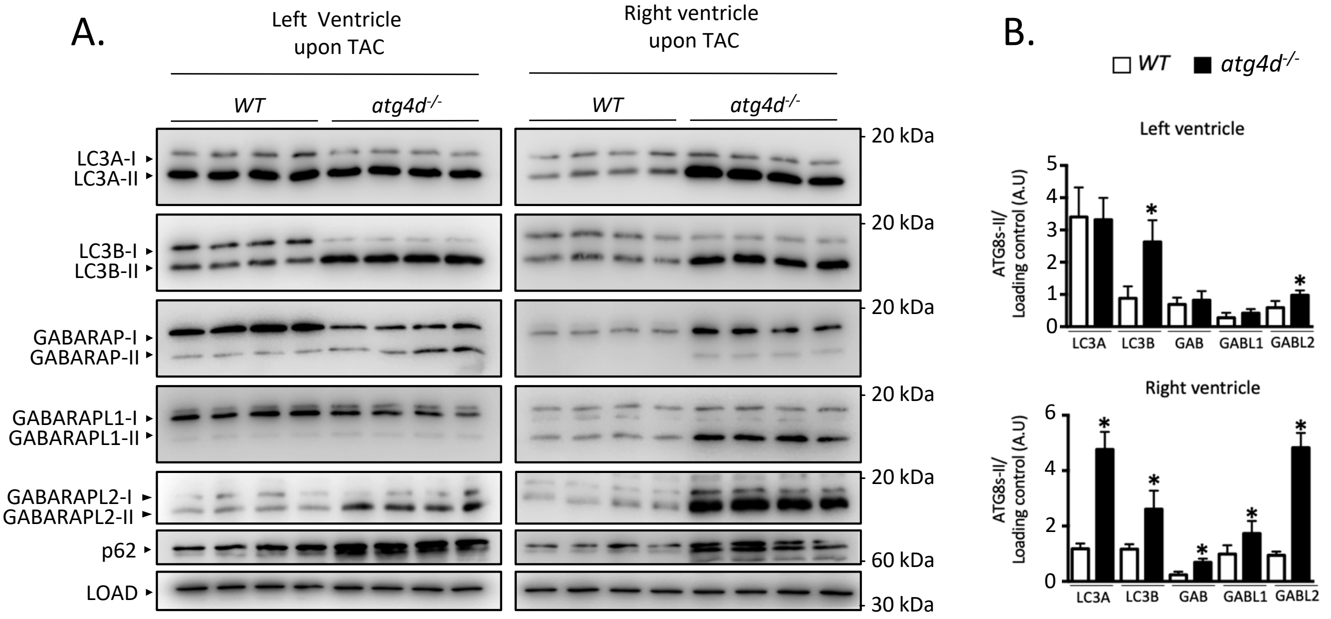

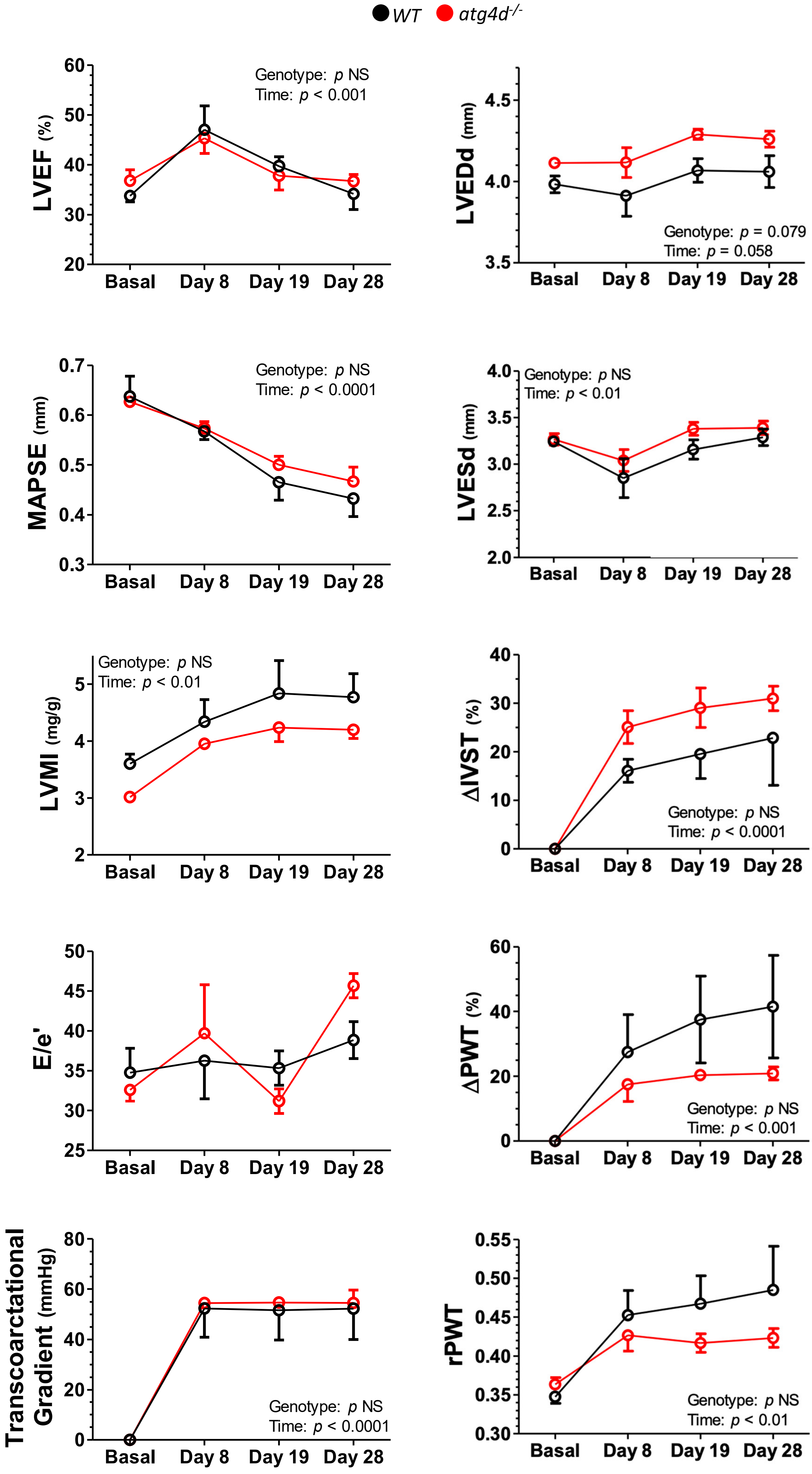

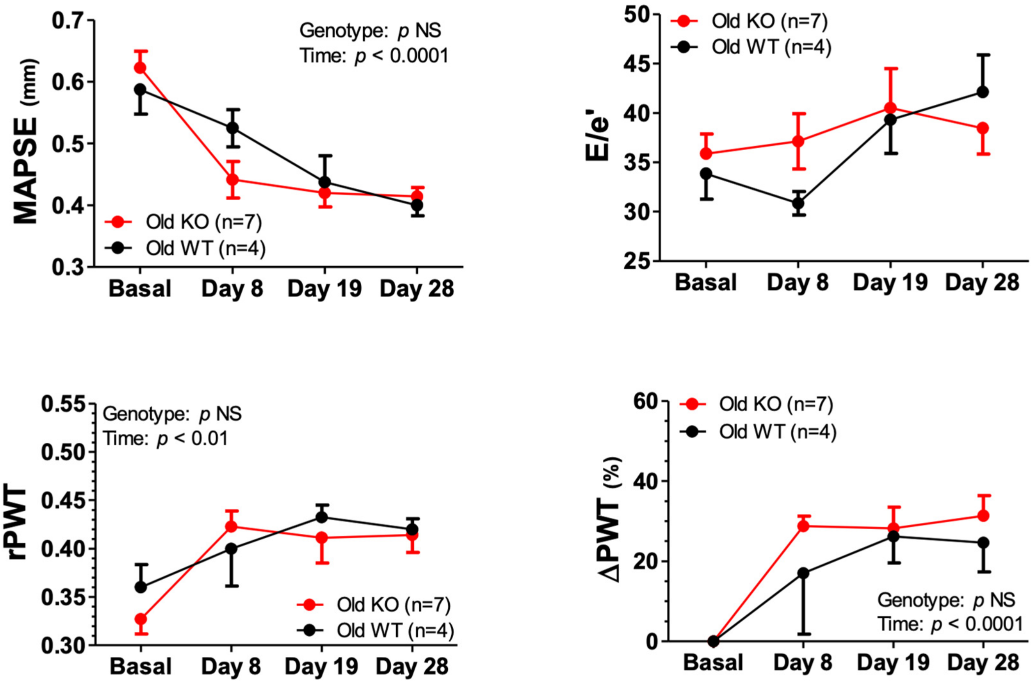
